# The role of the NLRP3 inflammasome in a CLN2 mouse model of neuroinflammation

**DOI:** 10.64898/2026.10.12.758052

**Authors:** Kathryn E. Sánchez, Samantha Elizalde, Nancy B. Schwartz, Miriam S. Domowicz

## Abstract

Neuronal ceroid lipofuscinoses, or Batten disease, comprises a group of inherited pediatric neurodegenerative disorders, and is characterized by the lysosomal accumulation of the fluorescent storage material lipofuscin. The human classical late-infantile form of the disorder (CLN2) is caused by mutations in the *CLN2*/*Tripeptidyl peptidase 1* (*TPP1*) gene. Comparable to human CLN2, a knockout of the *Tpp1* gene (*Tpp1 -/-*) in the mouse leads to progressive vision loss, ataxia, motor atrophy, seizures, and premature death. While we previously showed using bulk RNA-seq analysis that reactive gliosis develops between 3 and 4 months in this animal model, the role of the inflammasome in CLN2 remains unexplored. The inflammasome is a multiprotein complex formed through pattern recognition receptor engagement, which culminate in the cleavage and release of interleukin-1 beta. Here we report that while some inflammasome-associated genes are upregulated as early as 1-month in the *Tpp1 -/-* cerebellum, it is not until 3-months-old that all gene transcripts necessary for inflammasome assembly and function are upregulated, leading to IL1β release. In *Tpp1 -/-* slice cultures, we demonstrated that MCC950, an inhibitor of inflammasome assembly, downregulates transcriptional expression of genes associated with reactive microglia and inflammasome function. Furthermore, prophylactic administration of MCC950 in drinking water of *Tpp1 -/-* animals for two months is sufficient to lower the expression of these same genes in the forebrain and cerebellum, suggesting that inflammasome inhibitors are promising therapeutics for CLN2 and neurodegeneration broadly.

**Graphical Abstract:** The role of the NLRP3 inflammasome in a CLN2 model of neuroinflammation. The NLRP3 inflammasome is upregulated and assembled in *Tpp1-/-* microglia, leading to IL1β release. Treatment with an NLRP3 inflammasome inhibitor MCC950, in *ex-vivo* and *in-vivo* murine models of CLN2 neuroinflammation, results in the downregulation of inflammasome-associated genes.

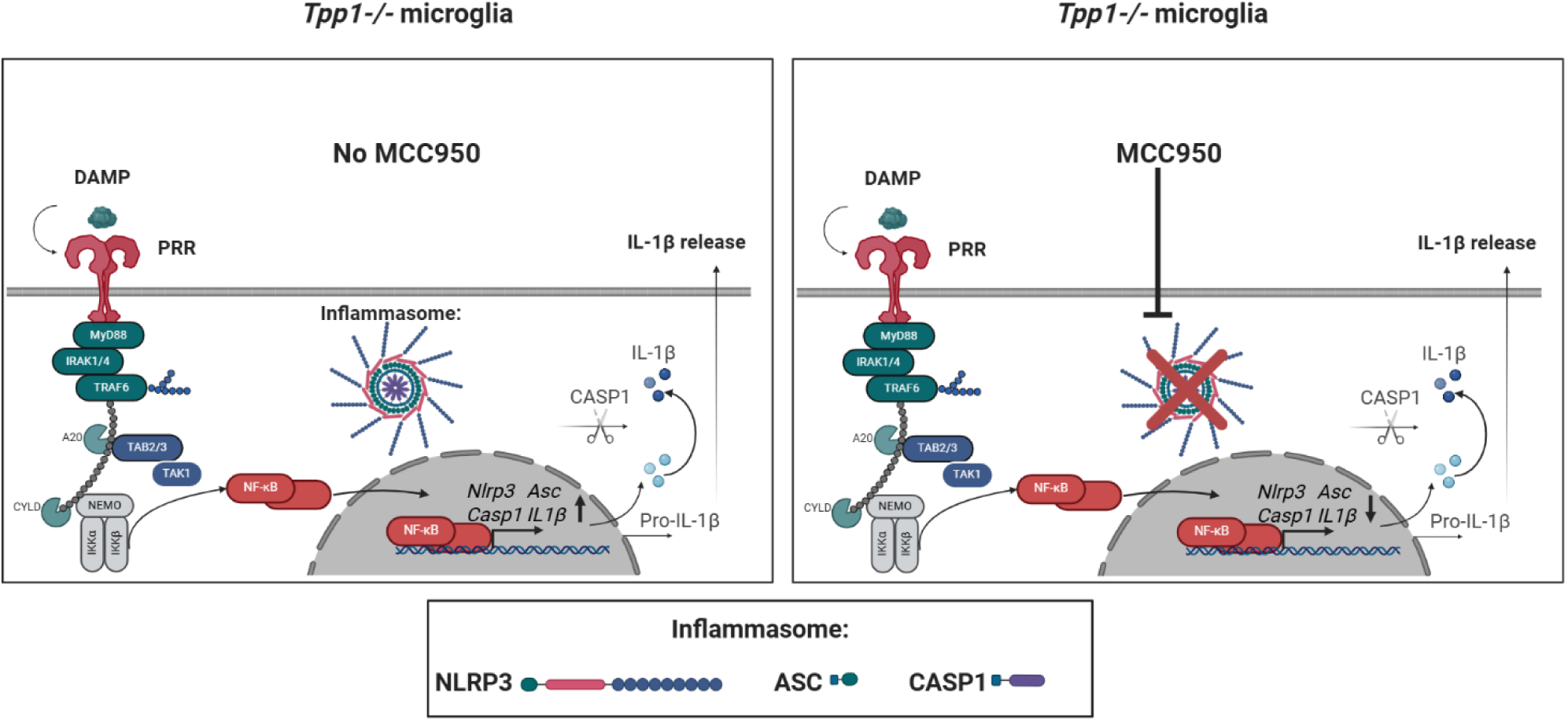

## Background

Neuronal Ceroid Lipofuscinoses (NCLs or Batten disease) are a group of pediatric, fatal neurodegenerative diseases characterized by lysosomal accumulation of autofluorescent storage material (ceroid lipofuscin). NCLs are classified as rare diseases and are estimated to occur in 2-4 out of every 100,000 live births (Mink et al. 2013; Kollmann et al. 2013). Specific NCL subtypes are caused by mutations in more than a dozen genes and have different ages of onset. In particular, classical late-infantile NCL (cLINCL; CLN2) in humans is due to specific mutations in the *Cln2/Tpp1* gene, which encodes the lysosomal enzyme tripeptidyl-peptidase 1(Sleat et al. 2004). CLN2 is typically diagnosed after the onset of seizures at 2-4 years of age. Due to severe neurodegeneration and limited treatment options, most patients die between 7 and 15 years old.

Tripeptidyl peptidase 1 deficient (*Tpp1 -/-*) mice display pathology similar to human cLINCL, making it an optimal model to study disease progression (Sleat et al. 2004). For example, these mice develop lipofuscin deposits, have profound neurodegeneration in the thalamus and cerebellum (Sleat et al. 2004; Domowicz et al. 2019, 2021), and die prematurely between 15 and 17 weeks of age (Sleat et al. 2004). We have documented that a key component of CLN2 pathology is the inflammatory response facilitated by microglia, the innate immune cells of the brain (Domowicz et al. 2019, 2021). Our prior studies have shown that microglia-related inflammatory genes *Cathepsin S* (*Ctss)* and *Cluster of Differentiation 68* (*Cd68*) are upregulated in *Tpp1 -/-* mouse brain at the late stage of disease (3-4 months of age) (Domowicz et al. 2019, 2021). However, the molecular pathways that drive the shift in microglia phenotype between 2- and 3-months of age in the *Tpp1-/-* model remain unknown. One such pathway is the NOD-like receptor pyrin domain containing 3 (NLRP3) inflammasome, which is elevated in the early stages of several neurodegenerative diseases (Codolo et al. 2013; White et al. 2017; Nazir et al. 2017; Panicker et al. 2019; Scott et al. 2020).

The inflammasome is a multiprotein complex produced by microglia that is assembled in two steps in response to aggregated proteins (danger-associated molecular patterns; DAMPs) (Venegas et al. 2017; Scott et al. 2020; Hulse and Bhaskar 2022). Priming occurs when a pattern recognition receptor (PRR) such as Toll-like receptor 4 (TLR 4) senses misfolded protein aggregates on the cell membrane, stimulating a protein kinase cascade that leads to the nuclear translocation of Nuclear Factor kappa-light-chain-enhancer of activated B cells (NFκB), which acts as a transcription factor to upregulate proinflammatory genes including *Interleukin-1 Beta* (*Il1b)* (Latz et al. 2013; Caplan and Maguire-Zeiss 2018; Hulse and Bhaskar 2022). Aggregated proteins are then phagocytosed leading to the activation of the inflammasome multiprotein complex consisting of NLRP3, ASC (Apoptosis-associated speck-like protein containing a Caspase Activation and Recruitment Domain), and Caspase-1 (Casp-1), followed by the cleavage of the inflammatory cytokine IL1β to promote its maturation (Hulse and Bhaskar 2022).

In other neurodegenerative disorders, proteins associated with inflammasome function are increased compared to controls. For example, IL1β and ASC are elevated in serum (Scott et al. 2020; Erhardt et al. 2021) and cerebrospinal fluid (Sánchez et al. 2022) of individuals with Alzheimer’s disease and related dementias (ADRD). IL1β and NLRP3 levels are also significantly increased in the plasma of individuals with Parkinson’s disease (PD)(Fan et al. 2020; Anderson et al. 2021), and chronic NLRP3 inflammasome activation is triggered by pathogenic misfolded α-synuclein (Codolo et al. 2013). The inflammasome has been investigated in the CLN3 mouse model of Batten Disease, *Cln3^Δex7/8^*, where a proinflammatory phenotype in *Cln3^Δex7/8^* microglia has been established due to IL1β release (Burkovetskaya et al. 2019). Treatment with ceramide, a lipid intermediate present in neurons and elevated in the CLN3 brain, led to the release of IL1β by *Cln3^Δex7/8^* microglia in culture. The role of the inflammasome was confirmed *in vivo* by crossing Cln3^Δex7/8^ and *Casp-1*-deficient mice to create Cln3^Δex7/8^ /*Casp-1^-/-^*animals (Burkovetskaya et al. 2019). Without *Casp-1*, *Cln3^Δex7/8^* line no longer displayed motor disfunction and astrocyte activation, emphasizing the importance of astrocytes and *Casp-1* in the disease process.

In addition to genetic interventions, pharmacological studies have identified an inhibitor of NLRP3 assembly, MCC950 (Dempsey et al. 2017; Perera et al. 2018; Ward et al. 2019; Jiao et al. 2020; Li et al. 2022; Liu et al. 2022; Naeem et al. 2024), thus providing a potential approach to prevent the release of IL1β downstream of the inflammasome (Dempsey et al. 2017; Jiao et al. 2020; Li et al. 2022; Hong et al. 2024; Naeem et al. 2024). For example, the administration of MCC950 in rodent models of spinal cord injury (Jiao et al. 2020), ADRD (Dempsey et al. 2017; Naeem et al. 2024), and Huntington’s disease(Chen et al. 2022) resulted in reduction in inflammasome proteins such as NLRP3 and ASC, decrease in the buildup of pathological aggregates such as amyloid beta (Dempsey et al. 2017), and improvement in motor movement (Chen et al. 2022) and memory (Dempsey et al. 2017).

Here we report for the first time that some genes associated with the NLRP3 inflammasome (*Asc* and *Nlrp3*) are transcriptionally upregulated as early as 1-month of age in *Tpp1 -/-* mice. However, it is not until the symptomatic stage of disease at 3-months that all genes necessary for NLRP3 inflammasome formation (*Nlrp3*, *Asc*, *Casp-1*) and function (*Il1b*) are upregulated in the *Tpp1 -/-* cerebellum. We also provide evidence that both microglia and astrocytes produce ASC specks at this late stage of disease. Furthermore, in both a brain organotypic slice culture model and *in vivo Tpp1-/-* mice, we show that the NLRP3 specific inhibitor MCC950 interferes with the upregulation of genes. Lastly, these results emphasize the need to investigate the use of anti-inflammatory therapies in CLN2, providing the basis for future therapeutic interventions. More specifically, preventing inflammasome assembly may slow down disease progression and offer a supplement to current treatment paradigms, such as enzyme replacement therapy (Schulz et al. 2018; de Brito Sampaio et al. 2023), in CLN2. Overall, this study illustrates a potential therapeutic approach that could be applied to other genetic neurodegenerative disorders that involve neuroinflammation.

## Methods

### Animals

The use of mice for organotypic slice culture and all other animal work was performed at the University of Chicago where the protocol for all experiments with mice was approved by the Institutional Animal Care and Use Committee (IACUC). Mice were housed under 12-hour light/dark conditions and were administered chow and water *ad-libitum.* The mouse model for CLN2, in which *Tpp1* is disrupted by gene targeting, was a generous gift from Drs. P. Lobel and D.E. Sleat (Sleat et al. 2004) and has been previously described (Sleat et al. 2004; Domowicz et al. 2019, 2021). The line is referred to in this study as the *Tpp1 -/-* mutant and is maintained with *Tpp1* +/- heterozygous crosses. In all experiments, *Tpp1 -/-* homozygous mice were compared to wildtype or heterozygous (*Tpp1* +/- or *Tpp1* +/+) control siblings of matched age and sex, as indicated for both *in vivo* and *ex vivo* experiments. For MCC950 water administration, either water (Sham) or 0.3 mg/mL MCC950 was administered starting at 2 months of age for a duration of 8 weeks and sacrificed at 16 weeks of age. The same number of male and female mice were used for each experiment.

### Organotypic slice culture

The preparation for brain organotypic slices has previously been described (Polleux and Ghosh 2002; King et al. 2009). Briefly, hanging wells (Falcon; 353102) were coated with poly-L-lysine (Millipore; R2636) and laminin (Sigma; L2020) and incubated at 37°C overnight. After isolating the brain, the cerebellum and olfactory bulbs were removed, and the rest of the brain was mounted in 3% SeaPlaque Agarose (Lonza; 50100) in 1X HBSS Complete Media (1X HBSS, 2.5M HEPES, 30mM D-Glucose, 1mM CaCl_2_, 1mM MgSO_4,_ and 4 mM NaHCO_3_). Coronal brain slices of *Tpp1-/-* and control mice were collected with a Leica VT1000S vibratome. A minimum of 10 sections of 200 µM thickness from each brain per well, at a speed of 30.5 mm/s and frequency of 8 Hz, were collected in hanging wells (Falcon, 353102) and cultured on slice culture media (38% Basal Eagle Medium, 37% Complete Hanks’ Balanced Salt Solution (HBSS), 20mM D-Glucose, 1M L-glutamine, 100 U/mL Penicillin/0.1 mg/mL Streptomycin, and 24% Horse Serum) using 6-well plates (Falcon; 353502) and maintained at 37°C with 5% CO_2_. For MCC950 experiments, brain slice cultures were exposed to 10 or 50 nM MCC950 (APExBIO; B7946), or water (VEH) directly added to slice culture media for 6 hours (6h).

### Quantitative PCR

RNA was purified from Trizol-homogenized brain or slice cultures (Invitrogen; 5596018) using the Direct-zol™ RNA Miniprep Kit (Zymo Research; R2052) and reverse-transcribed with the High-Capacity Reverse Transcription Kit (Applied Biosystems; 4368814). RT-qPCRs were performed using the Sso Advanced TM Universal SYBR Green system (Bio-Rad Laboratories; 1725121) in the CFX Duet real time instrument (Bio-Rad Laboratories). Primers for *Gfap*, *Cd68*, and *Ctss* have been previously described (Domowicz et al. 2019). PrimeTime™ qPCR Primers (IDT) were utilized for *Il1b* (Mm.PT.58.41616450), *Asc* (Mm.PT.56a.42872867), *Nlrp3* (Mm.PT.58.6779853.gs), and *Casp-1* (Mm.PT.58.13005595). Relative normalized expression values were determined using the ΔΔCt methods (Pfaffl 2001) for the indicated target genes relative to *Ppia* (Peptidylprolyl isomerase A) in the control sample. The data are expressed as means ± standard deviations. Determinations were performed in triplicate and experiments were repeated with independent RNA samples at least 3 times with consistent results. To compare two groups, statistical significances were evaluated by the paired-samples Student’s t-test using the CSF-96 BioRad Maestro software package. For three or more groups, a One-Way Analysis of Variance (ANOVA) was used to compare *Tpp1-/-* samples with and without MCC950 treatment using GraphPad Prism, Version 8. Control *Tpp1 +/-* samples were used to normalize gene expression in all experiments. Values of p < 0.05 for the null hypothesis were considered significant.

### Immunohistochemistry (IHC)

Control and *Tpp1* -/- 4-month brains were fixed in 4% paraformaldehyde and processed for immunohistochemistry. Sections were blocked with 2% bovine serum albumin (Bovine serum albumin; Fisher, BP9706-100) for 2h and incubated with a 1:200 dilution of anti-rabbit ASC (RRID:AB_2490442; Adipogen; AG-25B-0006-C200), 1:500 anti-rat CD68 (RRID:AB_869007; Abcam; ab53444), or 1:500 anti-mouse GFAP (RRID: AB_477010; Sigma; G3893), overnight at 4°C. Alexa Fluor 488-conjugated anti-rabbit IgG (RRID: AB_143165; Invitrogen; A11008), Rhodamine Red X-conjugated mouse IgG (RRID: AB_2556550; Invitrogen; R6393), or Alexa Fluor 546-conjugated anti-rat IgG (RRID: AB_141738; Invitrogen; A11081) was used as secondary antibody and slices were counterstained with 4’,6-diamidino-2-phenylindole (DAPI; Invitrogen; D3571). Representative images from the cerebellum were obtained with a Leica SP5 confocal microscope with a minimum of three independent mice per genotype. Imaging was performed at the University of Chicago Integrated Light Microscopy Core (RRID: SCR_019197).

### Enzyme-Linked Immunosorbent Assay (ELISA)

The protein levels of mouse IL1β were determined with enzyme-linked immunosorbent assays (ELISAs) according to the manufacturer’s instructions (R&D Systems; DY401-05). Cerebellum and cortical tissue were manually homogenized in Tissue Protein Extraction Reagent (TPER; Thermo; 78510) and spun down for 5 min at 10,000 rpm to collect protein supernatant. 100μL of supernatant (450-800 mg protein) was placed into each well, which were pre-coated with capture antibody. After a 2h incubation, washes were performed followed by the addition of an enzyme-linked polyclonal antibody specific to the mouse antigen. After a 2 h incubation, washes were conducted again and subsequently incubated with substrate solution for 30 min and immediately read after the addition of stop solution with the Victor 3 microplate reader (Perkin Elmer). IL1β release was normalized to amount of protein in each sample determined by BCA assay (Pierce; 23225).

### Western blot

Cerebellum lysates from 4-month-old mice were manually homogenized in RIPA buffer and spun down for 5 min at 10,000 rpm to collect protein supernatant. Total protein was determined using BCA assay (Pierce; 23225) and equal amounts of protein were used for each sample.

Samples were run on 10% SDS-PAGE gels and transferred to nitrocellulose membrane (Millipore; HATF85R) at 50 V overnight. Primary antibodies used are anti-β-actin (mouse IgG, 1:1000, RRID: AB_476744 Sigma; A5441) and anti-NLRP3 (rabbit IgG, 1:1000, RRID:AB_10646484; Proteintech; 19771). Secondary antibodies used were HRP-conjugated rabbit anti-mouse IgG (RRID: AB_228427; Pierce; 31450) or goat anti-rabbit (RRID: AB_228341; Pierce; 31460), respectively. Signal was detected using the Clarity Western ECL Substrate (Biorad; 1705060S) and visualized with the Azure 300 Imager (Azure Biosystems). Raw data shown in Supplemental Figure 1.

### Statistical Analysis

For slice culture studies, one animal per genotype was collected. Each experiment was repeated a minimum of three times, with one representative experiment shown on the graph indicative of three qPCR technical triplicates. For MCC950 *in vivo* studies and ELISA data, 5 independent animals per group are shown on the graph. For all experiments, comparisons between two groups were evaluated for statistical significance using an unpaired Student’s t-test in GraphPad Prism (Version 8). For comparisons among three or more groups, statistical significance was evaluated using a one-way or two-way Analysis of Variance (ANOVA) followed by Tukey’s post-hoc test, using GraphPad Prism (Version 8).

## Results

### Astrocytes and microglia upregulate proinflammatory genes in *Tpp1 -/-* cerebellum

Gliosis of astrocytes and microglia is well documented in both neurodegenerative and lysosomal storage disorders (Aflaki et al. 2016a; Domowicz et al. 2019, 2021; Ługowska et al. 2019). In the *Tpp1-/-* model, we previously established that gliosis occurs in the late stage of disease in the cerebellum, a region associated with neurodegeneration and ataxia (Domowicz et al. 2019, 2021). However, to better understand the pathogenesis of CLN2, we further investigated how gene expression correlated with phagocytic microglia changes between the presymptomatic (1-2 months) and symptomatic stages of disease (3-4 months). To determine how inflammation drives early and late stages of disease, RNA was isolated from the cerebellum of *Tpp1-/-* mice at 1, 2, 3 and 4 months of age and lysosomal genes typically upregulated in phagocytic microglia such as *Cd68* and *Ctss* (Domowicz et al. 2019, 2021) were evaluated. At 2-months of age (8 weeks), *Cd68 and Ctss* were not upregulated in *Tpp1 -/-* mice compared to *Tpp1 +/-* controls (Figure 1A-1B), indicating that microglia have not yet developed a phagocytic phenotype during the early stages of disease. However, our analyses did confirm upregulation of these genes in 3 and 4-month-old *Tpp1 -/-* mice as previously reported (Domowicz et al. 2019, 2021), verifying the shift in microglia phenotype that occurs at the same age that symptoms begin to occur (3-months; Figure 1). Furthermore, the expression of these genes continues to increase and peaks at terminal stages of disease (4-months of age). *Glial Fibrillary Acidic Protein* (*Gfap*; Figure 1A), which is associated with astrogliosis, was also not upregulated until 3 months of age, and peaks in expression at 4-months of age as previously reported (Domowicz et al. 2019, 2021).

**Fig 1.**
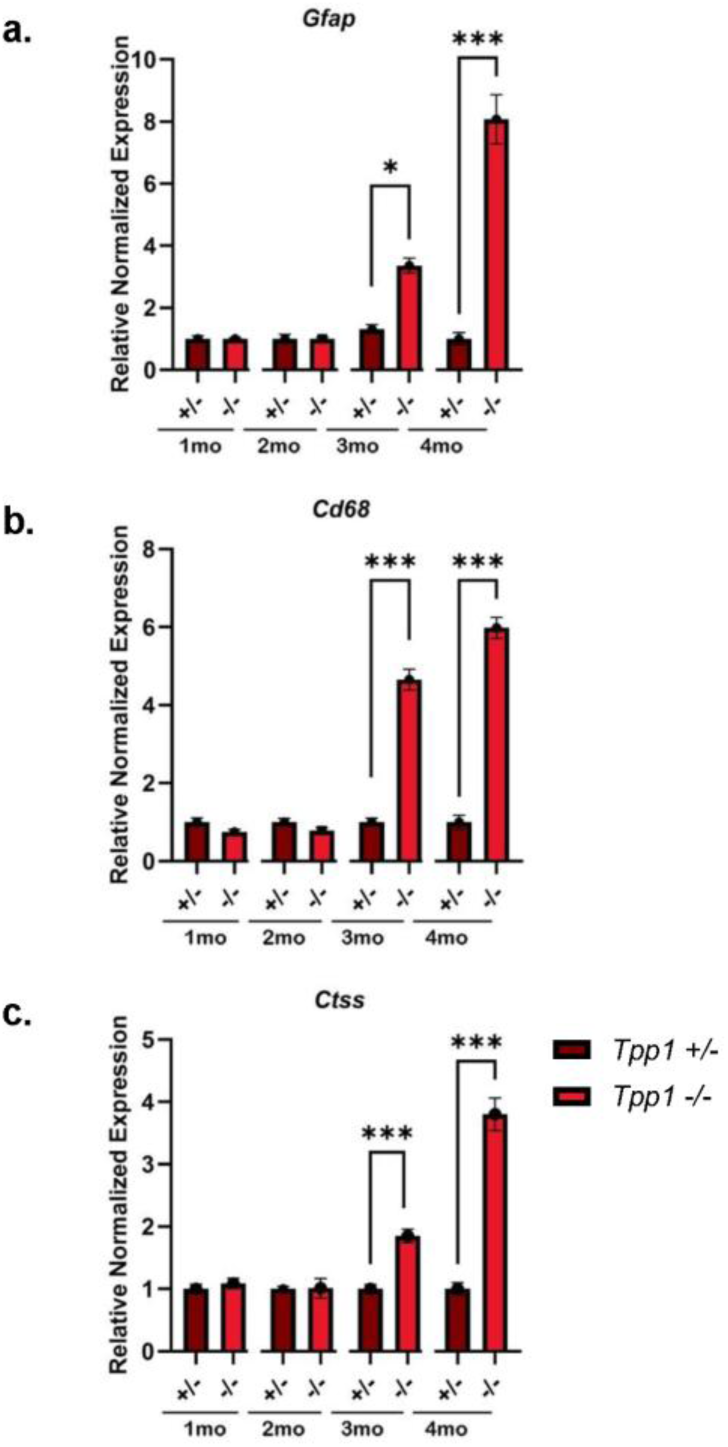
Genes associated with reactive glia are upregulated in the Tpp1 -/- mouse cerebellum at 3-4 months. RNA was isolated from the cerebellum of *Tpp1 -/-* and control mice at 1,2,3 and 4-months of age and qPCR of the indicated target genes was subsequently performed. Normalized expression of target genes was calculated relative to *Ppia* levels (housekeeping gene) with the ΔΔ CT method using BioRad CFX Maestro Manager. N=3 technical replicates. Values are reported as mean ± SEM; Student t-test compared to wildtype control of appropriate age; *P ≤ 0.05 and ***P ≤ 0.001. Figure representative of 3 independent experiments

### Inflammasome related genes are upregulated in *Tpp1 -/-* mice as early as 1-month of age

Knowing that microglia display an inflammatory phenotype in the *Tpp1 -/-* mouse model and that the inflammasome complex drives inflammation in other neurodegenerative disorders, we evaluated how inflammasome-related genes change over time in the *Tpp1 -/-* mouse using qRT-PCR at four different ages associated with distinct stages of disease progression. *Tpp1 -/-* mice expressed a statistically significant upregulation in genes associated with the inflammasome including *Asc* and *Nlrp3* as early as 1-month compared to controls (*Tpp1 +/-*), despite the lack of macrophagic microglia (Figure 2B-2C). In contrast, genes required for NLRP3 inflammasome assembly and function including *Nlrp3*, *Asc*, *Casp-1*, and *Il1b* are all upregulated by 3-months (12 weeks) of age in the *Tpp1-/-* mice (Figure 2A-2D). Furthermore, we evaluated NLRP3 and IL1β protein levels in the *Tpp1-/-* cerebellum at 4 months. We found that NLRP3 protein is increased in the *Tpp1 -/-* cerebellum compared to *Tpp1 +/-* control (Figure 3a). Inflammasome function is also evident at this timepoint since IL1β secretion is also elevated in the *Tpp1 -/-* cerebellum (Figure 3b) and cortex (Figure 3c) compared to *Tpp1 +/-* control, confirming that the transcriptional changes in these genes do translate to increased expression and release of IL1β protein.

**Fig 2.**
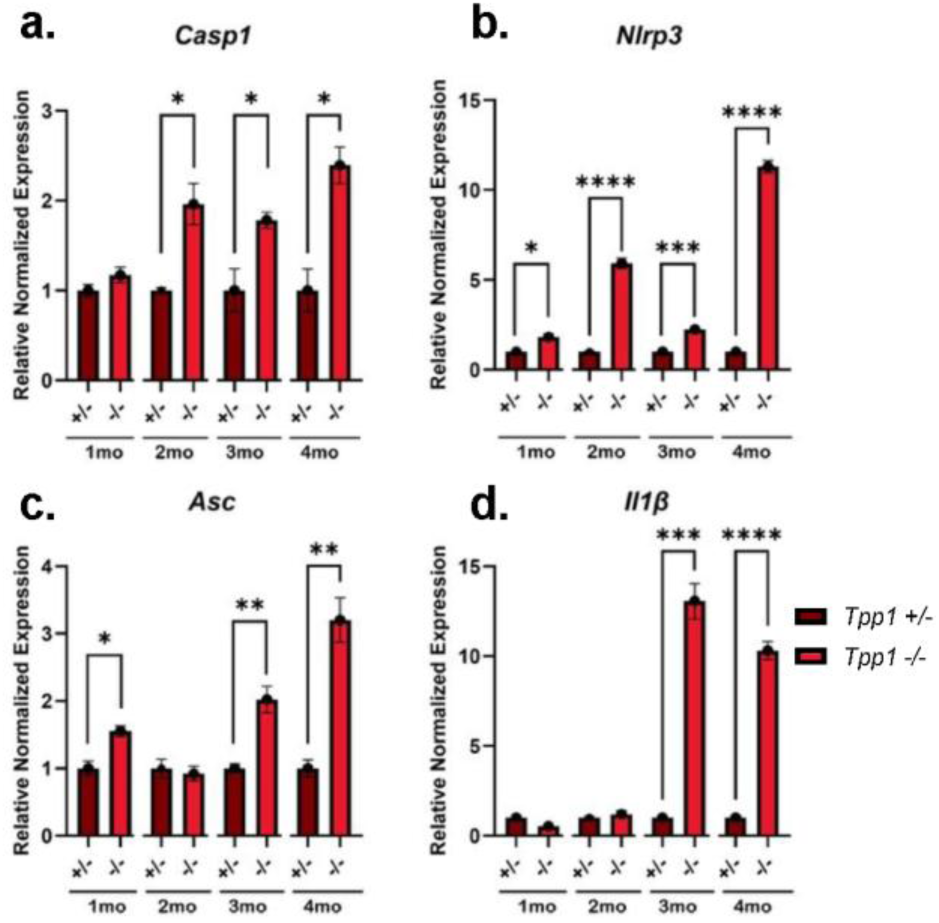
Genes associated with inflammasome assembly are upregulated at 1-month in the Tpp1 -/- mouse cerebellum. qPCR of the indicated target genes was performed with RNA extracted from the cerebellum of *Tpp1 -/-* and control mice at 1,2,3 and 4-months of age. Normalized expression of target genes was calculated relative to *Ppia* levels (housekeeping gene) with the ΔΔ CT method using BioRad CFX Maestro Manager. N=3 technical replicates. Values are reported as mean ± SEM; Student t-test compared to wildtype control of appropriate age; *P ≤ 0.05, ***P ≤ 0.001, and **** P ≤ 0.0001. Figure representative of 3 independent experiments. See Supplemental Figure 1 for raw data

**Fig 3.**
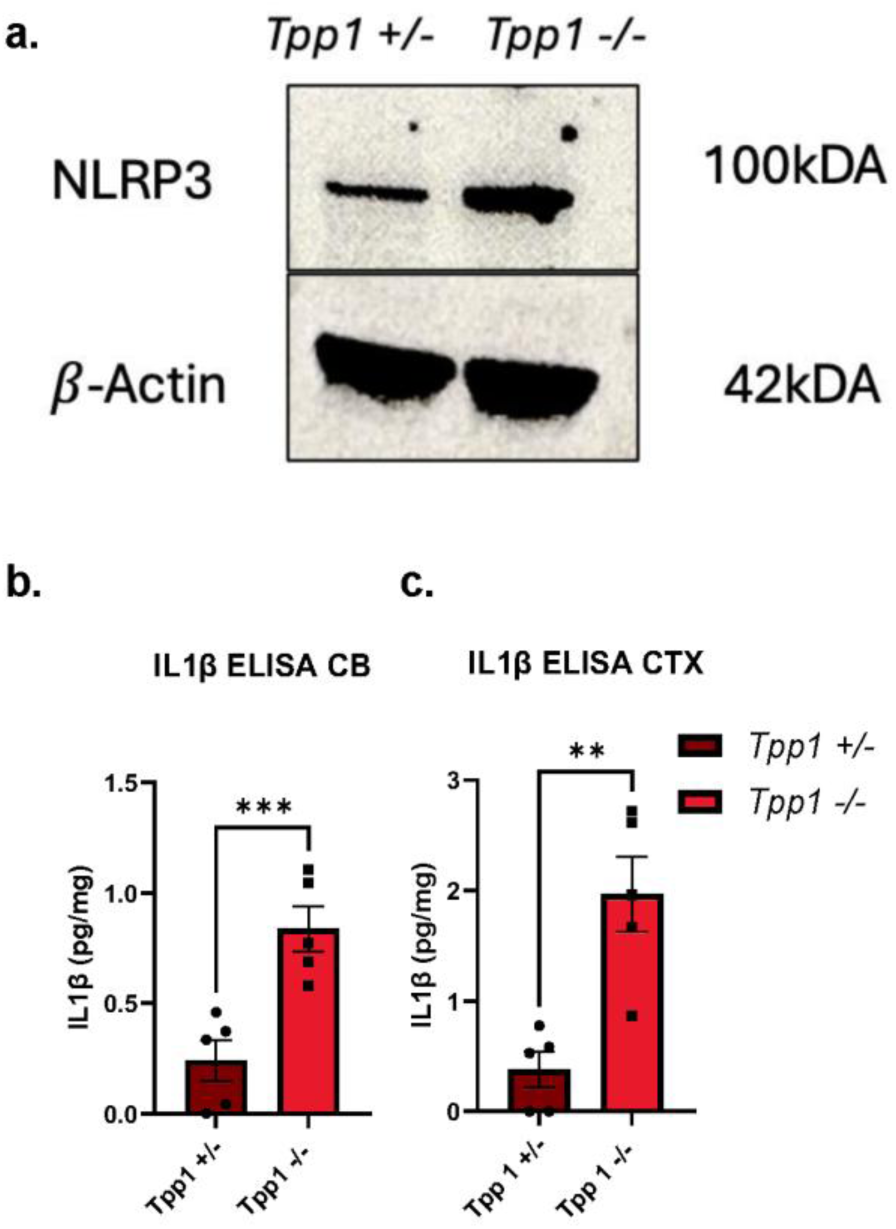
NLRP3 and IL1β are elevated in the Tpp1 -/- cerebellum at 4-months of age. Cerebellum was collected from 4-month-old *Tpp1 -/-* mice. NLRP3 levels were found to be elevated in *Tpp1 -/-*samples compared to *Tpp1 +/-* control homogenates by Western Blot (A). IL1β release was also elevated in *Tpp1-/-* cerebellum (B) and cortex (C) compared to controls as detected by ELISA. Values are reported as mean ± SEM; Student t-test compared to wildtype control of appropriate age; **P ≤ 0.01 and *** P ≤ 0.001

### Microglia and astrocytes express the ASC speck in *Tpp1 -/-* cerebellum

Due to the translational differences in IL1β that are evident in the *Tpp1 -/-* mouse, we next wanted to determine what cell type was responsible for its production. While previous investigations have established that the NLRP3 inflammasome is primarily expressed by microglia (Panicker et al. 2019; Jiang et al. 2021; Hulse and Bhaskar 2022), *Gfap* is also upregulated in *Tpp1 -/-* cerebellum at 4 months (Figure 1A). Therefore, to verify the cell type responsible for inflammasome assembly CLN2, IHC was conducted to visualize ASC specks in specific cell types. While ASC staining in the *Tpp1 +/-* control was not detectable, ASC specks were visible in the *Tpp1 -/-* mouse cerebellum at 4-months and primarily expressed by CD68^+^ microglia (Figure 4a) as well as some GFAP^+^ astrocytes as previously reported in other inflammatory conditions (Minkiewicz et al. 2013; Freeman et al. 2017) (Figure 4b), emphasizing the importance of glial cells in driving inflammasome assembly.

**Fig 4.**
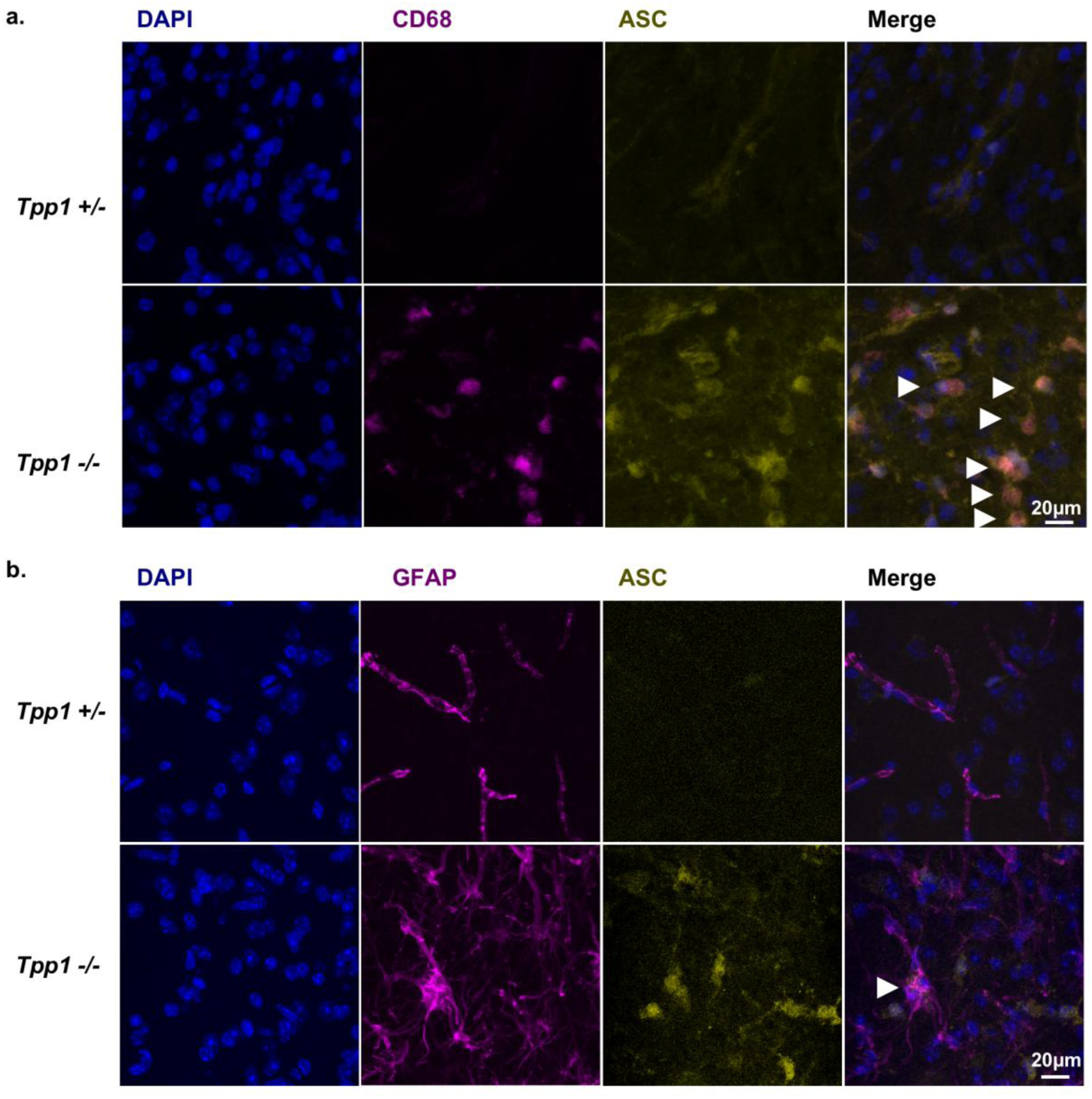
Inflammasome assembly occurs in microglia and astrocytes in the Tpp1 -/- cerebellum. Immunofluorescence was conducted in cerebellar sections of *Tpp1 -/-* and control mice at 4-months of age. Representative images show that ASC specks (yellow) colocalize (white arrow) with microglia (A; CD68) and astrocyte (B; GFAP) markers in the *Tpp1 -/-* cerebellum. N=3 biological replicates

### MCC950 treatment lowers central nervous system neuroinflammation in *Tpp1-/-* mice

Since inflammasome-mediated microglia inflammation is present in the *Tpp1 -/-* mice at 4-months of age, we evaluated the impact of NLRP3 inhibition and identified the role of NLRP3 specifically using the inhibitor of NLRP3 inflammasome assembly, MCC950 (Dempsey et al. 2017; Perera et al. 2018; Ward et al. 2019; Jiao et al. 2020; Li et al. 2022; Liu et al. 2022; Hong et al. 2024; Naeem et al. 2024). To test its therapeutic potential in the context of *Tpp1 -/-* pathology, we treated 4-month-old brain organotypic *ex vivo* slice cultures with MCC950 to evaluate proinflammatory gene expression using qPCR. After establishing effective dosages with 10µM and 50µM of MCC950 for 6h, it was observed that MCC950 reduced the expression of genes associated with phagocytosis (*Cd68;* Figure 5a) and inflammasome activation (*Asc*, *Il1b* and *Nlrp3*; Figure 5b-d) in the *Tpp1 -/-* slices at the 50µM dose specifically. Due to its effectiveness in the slice culture paradigm, these studies were expanded to the *in vivo* model by administering the drug to mice at 2-months of age prophylactically through drinking water (0.3 mg/mL; Figure 6a) until they were sacrificed at 4-months. It was found that while MCC950 did not significantly decrease the expression of *Cd68 and Ctss* to wildtype levels in the cerebellum (Figure 6c-6d) and forebrain (Figure 6h-6i), modest decreases were observed*. Gfap* expression was significantly elevated compared to wildtype levels in the cerebellum of *Tpp1 -/-* mice, with a significant decrease in MCC950 treated *Tpp1 -/-* mice (Figure 6b). GFAP was also lowered in the forebrain (Figure 6g), suggesting that MCC950 may directly or indirectly reduce astrogliosis (Domowicz et al. 2019, 2021). While *Nlrp3* expression was not changed in the cerebellum (Figure 6e), it was lowered in the forebrain (Figure 6j) after MCC950 administration likely due to more profound neurodegeneration in this region (Sleat et al. 2004). However, *Il1b* expression was lowered in both the cerebellum (Figure 6f) and forebrain (Figure 6k), confirming that MCC950 specifically decreases the expression of this important downstream target.

**Fig 5.**
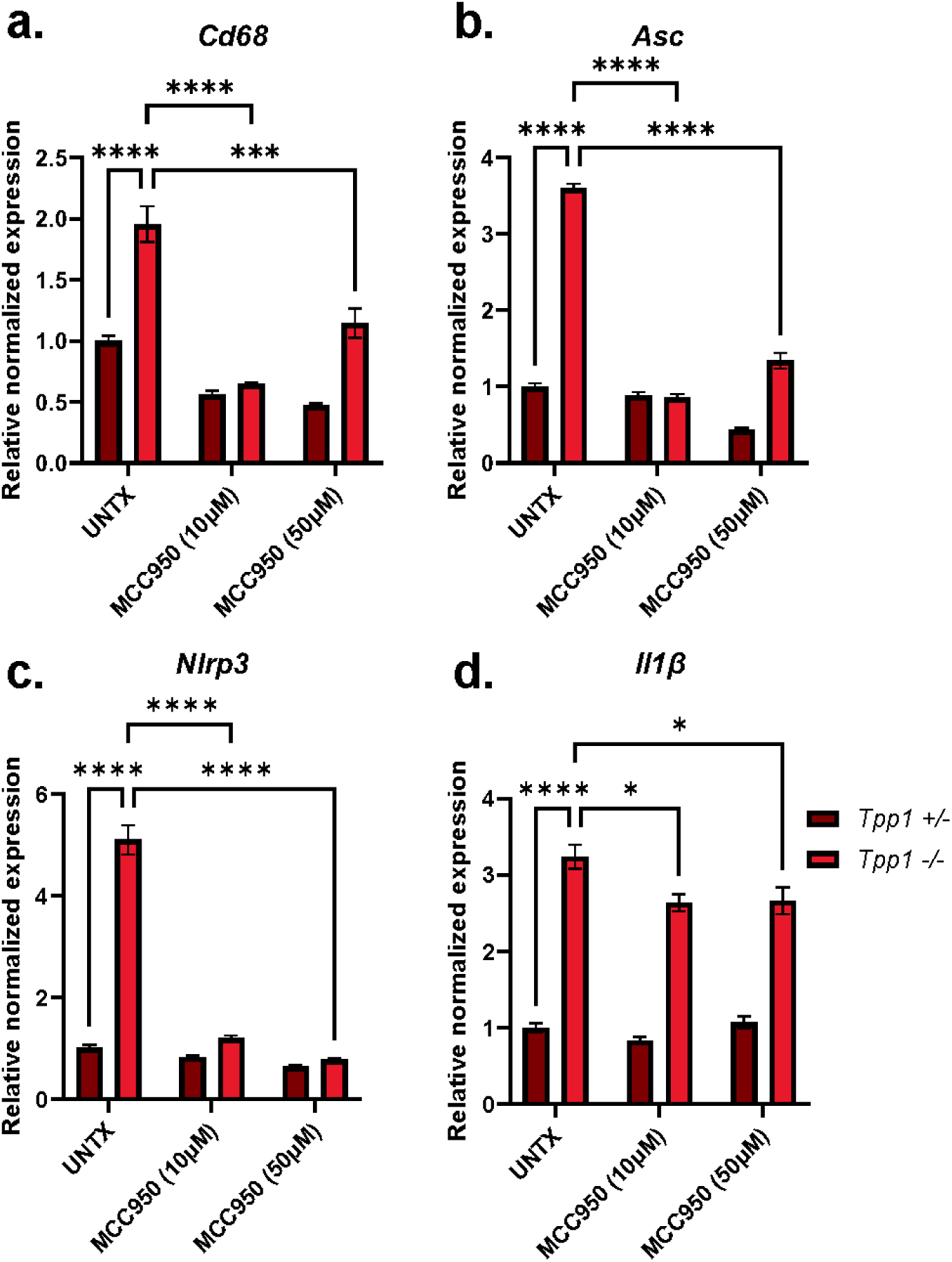
Upregulated genes associated with inflammasome assembly are inhibited by MCC950 in ex vivo organotypic slices cultures. Vibratome brain slices from 4-months in *Tpp1 -/-* and controls were collected. After 24-h in culture, slices were treated for 6h with MCC950 (10 and 50μM) and collected for RNA extraction to conduct qPCR. Normalized expression of target genes was calculated relative to *Ppia* levels (housekeeping gene) with the ΔΔCT method using BioRad CFX Maestro Manager. N=3 technical replicates. Values are reported as mean ± SEM; One-way ANOVA comparison with Tukey post hoc correction. While all groups were compared to each other, significance is shown to emphasize differences in *Tpp1-/-* in response to MCC950; ns= no significance, *P ≤ 0.05, ***P ≤ 0.001, and **** P ≤ 0.0001. Figure representative of 3 independent experiments

**Fig 6.**
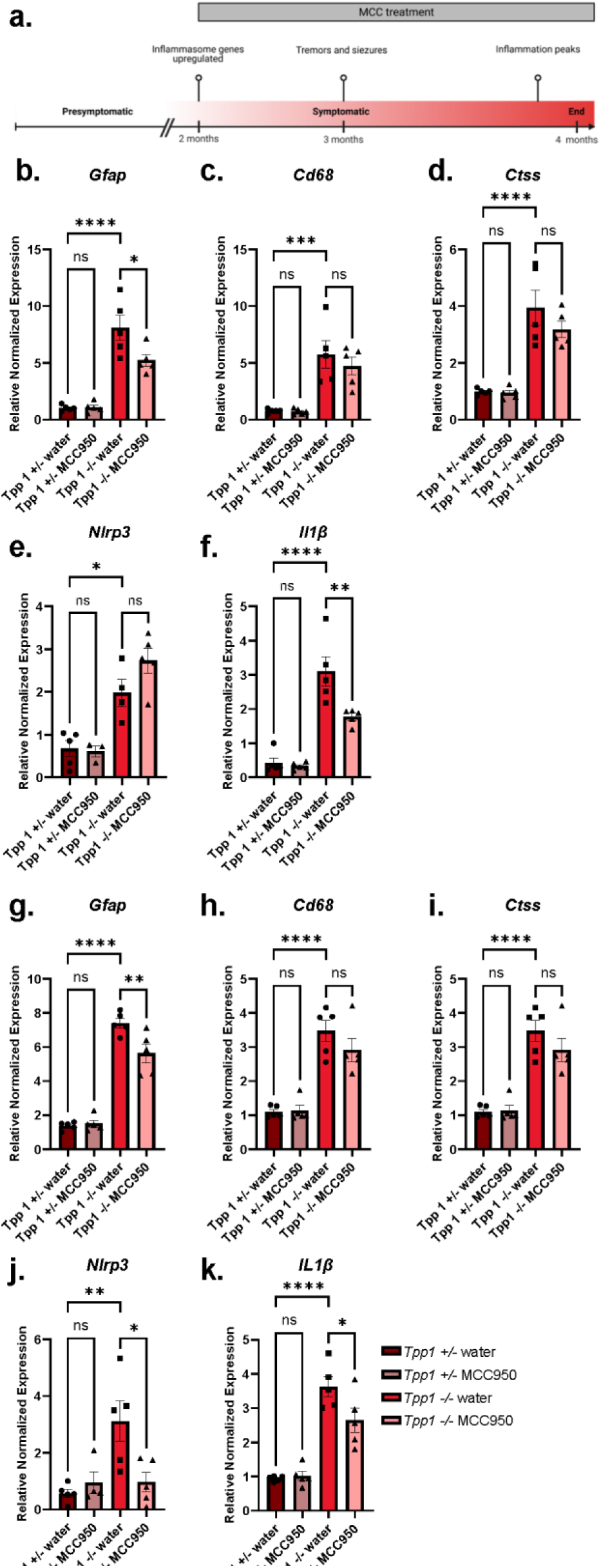
MCC950 inhibits genes associated with astrocytes and microglia activation when directly administered to Tpp1 -/- mice. MCC950 was administered to *Tpp1-/-* mice for the duration of 2-months in their drinking water and compared to control shams after sacrifice at 4-months of age, a timeframe chosen due to the chronological development of symptoms and pathology of disease (6A; Created with BioRender). qPCR of target genes was conducted from samples in the cerebellum (6B-6F) and forebrain (6G-6K). Normalized expression of target genes was calculated relative to *Ppia* levels (housekeeping gene) with the ΔΔCT method using BioRad CFX Maestro Manager. N=3-4 biological replicates. Values are reported as mean ± SEM; One-way ANOVA with Tukey post hoc correction; ns= no significance, *P ≤ 0.05, **P ≤ 0.01, *** P ≤ 0.001, and *** P ≤ 0.001. Figure representative of 3 independent experiments

Due to its effectiveness in the slice culture paradigm, these studies were expanded to the *in vivo* model by administering the drug to mice at 2-months of age prophylactically through drinking water (0.3 mg/mL; Figure 6a) until they were sacrificed at 4-months. It was found that while MCC950 did not significantly decrease the expression of *Cd68 and Ctss* to wildtype levels in the cerebellum (Figure 6c-6d) and forebrain (Figure 6h-6i), modest decreases were observed*. Gfap* expression was significantly elevated compared to wildtype levels in the cerebellum of *Tpp1 -/-* mice, with a significant decrease in MCC950 treated *Tpp1 -/-* mice (Figure 6b). GFAP was also lowered in the forebrain (Figure 6g), suggesting that MCC950 may directly or indirectly reduce astrogliosis (Domowicz et al. 2019, 2021). While *Nlrp3* expression was not changed in the cerebellum (Figure 6e), it was lowered in the forebrain (Figure 6j) after MCC950 administration likely due to more profound neurodegeneration in this region (Sleat et al. 2004). However, *Il1b* expression was lowered in both the cerebellum (Figure 6f) and forebrain (Figure 6k), confirming that MCC950 specifically decreases the expression of this important downstream target.

## Discussion

Microglia facilitate neuroinflammation and drive disease progression as they respond to neuronal death and lysosomal dysfunction (Aflaki et al. 2016a; Burkovetskaya et al. 2019; Domowicz et al. 2019, 2021). Disease-associated microglia (DAM) in particular have been implicated in numerous neurodegenerative disorders including PD, ADRD, and neuronal ceroid lipofuscinosis (Xiong and Kielian 2013; Burkovetskaya et al. 2019; Domowicz et al. 2019, 2021). DAM are microglia that downregulate homeostatic genes (*P2ry12/P2ry13, Cx3cr1, Tmem119*) and upregulate genes associated with phagocytosis and lipid metabolism (*Apoe, Lpl, Cst7, Ctsd, Trem2, Tyrobp*) (Cheng and Ho 2025) and are detected in neurodegenerative disorders. While DAM is most commonly observed in the context of ADRD (Rangaraju et al. 2018) and PD (Samant et al. 2024), functions and pathways associated with DAM have not been studied extensively in CLN2. More recently, the role of microglia has become better understood in the animal models of CLN2. Bulk RNA sequencing(Domowicz et al. 2019, 2021) showed transcriptional upregulation of genes associated with microglial activation at 3 and 4-months of age in the cerebellum in the CLN2 mouse, a time in which clinical symptoms such as ataxia and seizures also emerge. However, other microglia pathways such as the inflammasome have not been examined in CLN2 despite its established role in other neurodegenerative diseases (Abulafia et al. 2009; Codolo et al. 2013; Latz et al. 2013; Aflaki et al. 2016b; Saresella et al. 2016; White et al. 2017; Nazir et al. 2017; Ismael et al. 2018; Ward et al. 2019; Fan et al. 2020; Jiao et al. 2020; Scott et al. 2020; Zhang et al. 2020; Anderson et al. 2021; Hulse and Bhaskar 2022). Nor is it known if inflammasome assembly drives the phenotypic shift in microglia observed in the *Tpp1 -/-* mouse model of CLN2 between 2- and 3-months of age, correlating with the onset of symptoms that also occur at this stage of disease progression.

To better characterize microglia pathways in the chronological context of CLN2, this study showed that classic proinflammatory genes are not upregulated before the initial onset of clinical symptoms such as seizures and ataxia. Our data also corroborates that specific brain regions like the cerebellum, relevant to pathology and ataxia, also demonstrate neuroinflammation (Domowicz et al. 2019, 2021). Although genes upregulated in phagocytic microglia such as *Cd68* and *Ctss* are not upregulated until 3-months of age in *Tpp1-/-* mice, proinflammatory genes associated with the inflammasome such as *Asc, Nlrp3* and *Casp-1* are upregulated as early as 1-month of age. However, it is not until 3-months of age that all genes associated with the NLRP3 inflammasome complex (*Nlrp3*, *Asc*, *Casp1*) and function (*Il1b*) are upregulated. Given this chronological timeline, these findings indicate that microglia contribute to the assembly of the NLRP3 inflammasome and chronic inflammation in CLN2. These results also suggest that ASC, NLRP3, and Casp-1 have unique functions in early stages of inflammation separate from IL1β processing. ASC alone contributes to caspase-independent cell death (Willingham et al. 2007; Huang et al. 2009; Duncan et al. 2009), adaptive immunity (Deng et al. 2019; Ivanov et al. 2020), and antigen presentation (Sahillioʇlu and Özören 2015). NLRP3 also has functions not associated with canonical inflammasome signaling such as Casp-1 ubiquitination (Van Opdenbosch et al. 2014a), transcriptional regulation of *Il-4*, *Il-13*, *Il1r1*, *Icos*, and *Maf* (Bruchard et al. 2015; Honda et al. 2023), Smad2/3 phosphorylation (TGF-β pathway) (Lech et al. 2015), DNA damage response (ATM kinase) (Bodnar-Wachtel et al. 2023), and mitochondrial function in the presence of reactive oxygen species (Kim et al. 2019). Casp-1 can also promote pryoptosis independently of other inflammasome components (Barker et al. 2011; de Souza et al. 2021) and process other proinflammatory cytokines (Reinke et al. 2020).

Our finding that *Casp-1* (Figure 2A), *Nlrp3* (Figure 2B), and *Asc* (Figure 2C) are upregulated before *Il1b* (Figure 2D) also implicate distinct transcriptional regulation of *Nlrp3, Casp-1,* and *Asc* earlier in the disease process compared to *Il1b*. More specifically, the expression of *Nlrp3*, *Casp-1,* and *Asc* increases at an earlier timepoint (1-2 months), only to decrease before peaking again at a later timepoint (3-4 months). In fact, the transcription factor IRF1 has been shown to regulate both *Nlrp3* and *Casp-1* (Kuriakose et al. 2018; Benaoudia et al. 2019; Paik et al. 2021), raising the possibility it may act as a transcription factor for inflammasome-related genes and consequently contribute to the shift in microglia phenotype that occurs between 2-months and 3-months of age. Thus, Interferon-gamma (IFNγ) may drive early inflammation through IRF1, since it regulates both *Nlrp3* and *Casp-1* (Kuriakose et al. 2018; Benaoudia et al. 2019; Paik et al. 2021), while the transcription factor NFκB may promote a shift to chronic microglial activation which also includes the upregulation of *Il1b* (Christgen et al. 2020) at 3- and 4-months in the *Tpp1 -/-* mouse. In the context of the inflammasome, this suggests that ASC independent *Nlrp3* upregulation is taking place pre-symptomatically. More importantly, this indicates that canonical inflammasome function, including IL1β secretion, likely only occurs once inflammation has emerged between 3 and 4-months of age (Cassel et al. 2014; Lukens et al. 2014; Van Opdenbosch et al. 2014b; Nagar et al. 2021). This inflammation is profound later in disease at the onset of symptoms, indicating that inflammation may occur concomitantly with pathological processes including neurodegeneration.

In addition to the upregulation of all NLRP3 inflammasome related genes at 4-months of age in the *Tpp1 -/-* mouse, the inflammasome is fully assembled as ASC specks in CD68+ microglia by this time. Interestingly, ASC specks were also present in a small subpopulation of astrocytes, supporting more recent findings that astrocytes can assemble different forms of inflammasomes and contribute to the pro-inflammatory states observed in neurological diseases such as depression, epilepsy, PD, AD, pain, ischemic damage, and others (Mata-Martínez et al. 2022). In fact, when *Casp-1* is knocked out in astrocytes, hippocampal plasticity is prevented (Zengeler et al. 2025). Furthermore, in pathological environments the activation of the inflammasome in astrocytes is stimulated through the engagement of a PRR and the NFκB pathway in a manner similar to microglia (Nourbakhsh et al. 2021). Since a small subset of astrocytes express ASC, our findings suggest that the role of the inflammasome in *Tpp1-/-* astrocytes should be investigated more thoroughly. For example, after treatment with MCC950 *in vivo*, we found that *Gfap* expression was decreased in both the forebrain and cerebellum (Figure 6) providing further evidence that NLRP3 may directly or indirectly impact astrocyte/microglia crosstalk.

This study has also expanded our knowledge on the use of anti-inflammatory therapies in Batten disease (Kauss et al. 2020). While genetic inhibition of the inflammasome yielded an improvement of motor function in a CLN3 mouse model by genetically crossing with *Casp-1* deficient mice (Burkovetskaya et al. 2019), we are the first to report findings related to pharmaceutically inhibiting NLRP3 in a Batten disease model. The effectiveness of MCC950 in the context of neurodegeneration was corroborated using *ex vivo* slice cultures, which establish the specificity of the drug and show its ability to down-regulate genes associated with phagocytosis and the inflammasome in microglia. These findings emphasize how lysosomal destabilization activates the inflammasome during phagocytosis (Okada et al. 2014; Katsnelson et al. 2016). We have also shown that MCC950 can down-regulate genes critical to the inflammasome such as *Il1b* and *Nlrp3* in the forebrain and cerebellum when administered *in vivo* to 4-month-old *Tpp1 -/-* mice. These studies will be expanded in the future in order to optimize the dose, frequency of administration, and time of initial MCC950 administration. Moreover, we established a critical role of NLRP3 during the preclinical stage of disease and its importance in shifting microglia from the acute inflammatory state to a more chronic phenotype (Lull and Block 2010; Perry and Teeling 2013; Ajami et al. 2018). However, IL1β release was not significantly lowered in either the cortex or cerebellum after MCC treatment (Supplemental Figure 2), indicating that NLRP3 independent IL1β production should be further studied in the future. Furthermore, we report a consistent 25% decrease in *Il1b* message with MCC950 in both the *ex vivo* and *in vivo* model. However, IL1β is still being translated at the protein level (Supplemental Figure 2). Therefore, IL1β production is likely taking place as well through the engagement of other sensor proteins in the presence of MCC950. Thus, future investigations should characterize the role of alternative sensor proteins such as AIM2, NLRP12, and NLRP6 the CLN2 mouse model.

In conclusion, our data establish the inflammasome as a putative therapeutic target in the *Tpp1* -/- mouse model and suggest that inflammasome assembly may be a driver of inflammation due to its early upregulation. These studies have also identified an anti-inflammatory therapy that controls chronic inflammation and improves synaptic function in a mouse model of CLN2. Thus, inflammasome inhibitors may be a critical supplemental treatment that can complement enzyme replacement therapy in the clinic.

## Declarations

## Acknowledgments

We would like to thank our colleagues in the Schwartz lab for their assistance with this project. We also thank Drs. P. Lobel and D.E. Sleat for gifting the *Tpp1 -/-* mice line.

## Author Contributions

KES, SE, and MSD, planned and performed all experiments. KS wrote the initial draft and NBS and MSD provided feedback on scientific interpretation and subsequent drafts. All authors contributed to the article and approved the submitted version.

## Ethics approval and consent to participate

All animal procedures were performed in strict accordance with the institutional guidelines for the care and use of experimental animals approved by the University of Chicago Institutional Animal Care and Use Committee.

## Consent for Publication

Not applicable.

## Availability of Data

The datasets used and/or analyzed during the current study are available from the corresponding author on reasonable request.

## List of Abbreviations

NCL: Neuronal Ceroid Lipofuscinoses
cLINCL: Classical late-infantile neuronal ceroid lipofuscinoses
CLN2: Neuronal Ceroid Lipofuscinosis Type 2
Tpp1: Tripeptidyl peptidase 1
DAMPs: Danger-associated molecular patterns
PRR: Pattern recognition receptor
NLRP3: NLR family pyrin domain containing 3 (or NOD-like receptor pyrin domain containing 3)
ASC: Apoptosis-associated speck-like protein containing a CARD (Caspase Activation and Recruitment Domain)
IL1β: Interleukin-1 Beta
ADRD: Alzheimer’s disease and related dementias
PD: Parkinson’s disease
CLN3: Neuronal Ceroid Lipofuscinosis Type 3
HBSS: Hanks’ Balanced Salt Solution
ANOVA: Analysis of variance
IHC: Immunohistochemistry
DAPI: 4’,6-diamidino-2-phenylindole
DAM: Disease-associated microglia

## References

Abulafia DP, De Rivero Vaccari JP, Lozano JD, et al (2009) Inhibition of the inflammasome complex reduces the inflammatory response after thromboembolic stroke in mice. Journal of Cerebral Blood Flow and Metabolism 29:534–544. 10.1038/jcbfm.2008.143

Aflaki E, Moaven N, Borger DK, et al (2016a) Lysosomal storage and impaired autophagy lead to inflammasome activation in Gaucher macrophages. Aging Cell 15:77–88. 10.1111/ACEL.12409

Ajami B, Samusik N, Wieghofer P, et al (2018) Single-cell mass cytometry reveals distinct populations of brain myeloid cells in mouse neuroinflammation and neurodegeneration models. Nat Neurosci 21:541–551. 10.1038/S41593-018-0100-X

Anderson FL, von Herrmann KM, Andrew AS, et al (2021) Plasma-borne indicators of inflammasome activity in Parkinson’s disease patients. NPJ Parkinsons Dis 7:1–12. 10.1038/S41531-020-00147-6

Barker BR, Taxman DJ, Ting JPY (2011) Cross-regulation between the IL-1β/IL-18 processing inflammasome and other inflammatory cytokines. Curr Opin Immunol 23:591. 10.1016/J.COI.2011.07.005

Benaoudia S, Martin A, Gamez MP, et al (2019) A genome-wide screen identifies IRF2 as a key regulator of caspase-4 in human cells. EMBO Rep 20:e48235. 10.15252/EMBR.201948235

Bodnar-Wachtel M, Huber AL, Gorry J, et al (2023) Inflammasome-independent NLRP3 function enforces ATM activity in response to genotoxic stress. Life Sci Alliance 6:1–14. 10.26508/LSA.202201494

Bruchard M, Rebé C, Derangère V, et al (2015) The receptor NLRP3 is a transcriptional regulator of TH2 differentiation. Nat Immunol 16:859–870. 10.1038/NI.3202

Burkovetskaya M, Bosch ME, Karpuk N, et al (2019) Caspase 1 activity influences juvenile Batten disease (CLN3) pathogenesis. J Neurochem 148:652–668. 10.1111/JNC.14480

Caplan IF, Maguire-Zeiss KA (2018) Toll-like receptor 2 signaling and current approaches for therapeutic modulation in synucleinopathies. Front. Pharmacol 9:1–18 10.3389/fphar.2018.00417

Cassel SL, Janczy JR, Bing X, et al (2014) Inflammasome-independent IL-1β mediates autoinflammatory disease in Pstpip2-deficient mice. Proc Natl Acad Sci U S A 111:1072– 1077. 10.1073/PNAS.1318685111

Chen KP, Hua KF, Tsai FT, et al (2022) A selective inhibitor of the NLRP3 inflammasome as a potential therapeutic approach for neuroprotection in a transgenic mouse model of Huntington’s disease. J Neuroinflammation 19:1–17. 10.1186/s12974-022-02419-9

Cheng YH, Ho MS (2025) Disease-associated microglia in neurodegenerative diseases: Friend or foe? PLoS Biol 23:e3003426. 10.1371/JOURNAL.PBIO.3003426

Christgen S, Place DE, Kanneganti TD (2020) Toward targeting inflammasomes: insights into their regulation and activation. Cell Research 2020 30:4 30:315–327. 10.1038/s41422-020-0295-8

Codolo G, Plotegher N, Pozzobon T, et al (2013) Triggering of inflammasome by aggregated α-synuclein, an inflammatory response in synucleinopathies. PLoS One 8:e55375. 10.1371/journal.pone.0055375

de Brito Sampaio LP, de Manreza MLG, Pessoa A, et al (2023) Clinical management and diagnosis of CLN2 disease: consensus of the Brazilian experts group. Arq Neuropsiquiatr 81:284–295. 10.1055/S-0043-1761434

de Souza JG, Starobinas N, Ibañez OCM (2021) Unknown/enigmatic functions of extracellular ASC. Immunology 163:377–388. 10.1111/IMM.13375

Dempsey C, Rubio Araiz A, Bryson KJ, et al (2017) Inhibiting the NLRP3 inflammasome with MCC950 promotes non-phlogistic clearance of amyloid-β and cognitive function in APP/PS1 mice. Brain Behav Immun 61:306–316. 10.1016/j.bbi.2016.12.014

Deng J, Yu XQ, Wang PH (2019) Inflammasome activation and Th17 responses. Mol Immunol 107:142–164. 10.1016/j.molimm.2018.12.024

Domowicz MS, Chan WC, Claudio-Vázquez P, et al (2021) Brain transcriptome analysis of a CLN2 mouse model as a function of disease progression. J Neuroinflammation 18:262. 10.1186/S12974-021-02302-Z

Domowicz MS, Chan WC, Claudio-Vázquez P, et al (2019) Global brain transcriptome analysis of a Tpp1 Neuronal Ceroid Lipofuscinoses mouse model. ASN Neuro 11: 1759091419843393. 10.1177/1759091419843393

Duncan JA, Gao X, Huang MT-H, et al (2009) Neisseria gonorrhoeae activates the proteinase cathepsin B to mediate the signaling activities of the NLRP3 and ASC-containing inflammasome. J Immunol 182:6460–6469. 10.4049/JIMMUNOL.0802696

Erhardt EB, Adair JC, Knoefel JE, et al (2021) Inflammatory Biomarkers Aid in Diagnosis of Dementia. Front Aging Neurosci 13:717344. 10.3389/FNAGI.2021.717344

Fan Z, Pan YT, Zhang ZY, et al (2020) Systemic activation of NLRP3 inflammasome and plasma α-synuclein levels are correlated with motor severity and progression in Parkinson’s disease. J Neuroinflammation 17:1–11. 10.1186/S12974-019-1670-6

Freeman L, Guo H, David CN, et al (2017) NLR members NLRC4 and NLRP3 mediate sterile inflammasome activation in microglia and astrocytes. Journal of Experimental Medicine 214:1351–1370. 10.1084/jem.20150237

Honda TSB, Ku J, Anders HJ (2023) Cell type-specific roles of NLRP3, inflammasome-dependent and -independent, in host defense, sterile necroinflammation, tissue repair, and fibrosis. Front Immunol 14:1214289. 10.3389/FIMMU.2023.1214289

Hong Y, Wei C, Fu M, et al (2024) MCC950 alleviates seizure severity and angiogenesis by inhibiting NLRP3/ IL-1β signaling pathway-mediated pyroptosis in mouse model of epilepsy. Int Immunopharmacol 126: 111236. 10.1016/J.INTIMP.2023.111236

Huang MT-H, Taxman DJ, Holley-Guthrie EA, et al (2009) Critical role of apoptotic speck protein containing a caspase recruitment domain (ASC) and NLRP3 in causing necrosis and ASC speck formation induced by Porphyromonas gingivalis in human cells. J Immunol 182:2395–2404. 10.4049/JIMMUNOL.0800909

Hulse J, Bhaskar K (2022) Crosstalk Between the NLRP3 Inflammasome/ASC Speck and Amyloid Protein Aggregates Drives Disease Progression in Alzheimer’s and Parkinson’s Disease. Front Mol Neurosci 15: 805169. 10.3389/FNMOL.2022.805169

Ismael S, Zhao L, Nasoohi S, Ishrat T (2018) Inhibition of the NLRP3-inflammasome as a potential approach for neuroprotection after stroke. Sci Rep 8:1–9. 10.1038/s41598-018-24350-x

Ivanov K, Garanina E, Rizvanov A, Khaiboullina S (2020) Inflammasomes as Targets for Adjuvants. Pathogens 9:252. 10.3390/PATHOGENS9040252

Jiang S, Maphis NM, Binder J, et al (2021) Proteopathic tau primes and activates interleukin-1β via myeloid-cell-specific MyD88- and NLRP3-ASC-inflammasome pathway. Cell Rep 36:109720. 10.1016/J.CELREP.2021.109720

Jiao J, Zhao G, Wang Y, et al (2020) MCC950, a Selective Inhibitor of NLRP3 Inflammasome, Reduces the Inflammatory Response and Improves Neurological Outcomes in Mice Model of Spinal Cord Injury. Front Mol Biosci 7:519495. 10.3389/FMOLB.2020.00037

Katsnelson MA, Lozada-Soto KM, Russo HM, et al (2016) Cell Signaling: Proteins, Pathways and Mechanisms: NLRP3 inflammasome signaling is activated by low-level lysosome disruption but inhibited by extensive lysosome disruption: roles for K+ efflux and Ca2+ influx. Am J Physiol Cell Physiol 311:C83. 10.1152/AJPCELL.00298.2015

Kauss V, Dambrova M, Medina DL (2020) Pharmacological approaches to tackle NCLs. Biochimica et Biophysica Acta (BBA) - Molecular Basis of Disease 1866:165553. 10.1016/J.BBADIS.2019.165553

Kim YG, Kim SM, Kim KP, et al (2019) The role of inflammasome-dependent and inflammasome-independent nlrp3 in the kidney. Cells 8: 1389. 10.3390/CELLS8111389

King LA, Schwartz NB, Domowicz MS (2009) Glial migratory streams in the developing hindbrain: A slice culture approach. J Neurosci Methods 177:30–43. 10.1016/J.JNEUMETH.2008.09.019

Kollmann K, Uusi-Rauva K, Scifo E, et al (2013) Cell biology and function of neuronal ceroid lipofuscinosis-related proteins. Biochim Biophys Acta Mol Basis Dis 1832:1866–1881. 10.1016/j.bbadis.2013.01.019

Kuriakose T, Zheng M, Neale G, Kanneganti T-D (2018) IRF1 is a transcriptional regulator of ZBP1 promoting NLRP3 inflammasome activation and cell death during influenza virus infection. J Immunol 200:1489. 10.4049/JIMMUNOL.1701538

Latz E, Xiao TS, Stutz A (2013) Activation and regulation of the inflammasomes. Nat. Rev. Immunol. 13:397–411. 10.1038/nri3452

Lech M, Lorenz G, Kulkarni OP, et al (2015) NLRP3 and ASC suppress lupus-like autoimmunity by driving the immunosuppressive effects of TGF-β receptor signaling. Ann Rheum Dis 74:2224–2235. 10.1136/annrheumdis-2014-205496

Li H, Guan Y, Liang B, et al (2022) Therapeutic potential of MCC950, a specific inhibitor of NLRP3 inflammasome. Eur J Pharmacol 928:175091. 10.1016/J.EJPHAR.2022.175091

Liu Q, Zhang MM, Guo MX, et al (2022) Inhibition of microglial NLRP3 with MCC950 attenuates microglial morphology and NLRP3/Caspase-1/IL-1β signaling in stress-induced mice. Journal of Neuroimmune Pharmacology 17:503–514. 10.1007/S11481-021-10037-0

Ługowska A, Hetmańczyk-Sawicka K, Iwanicka-Nowicka R, et al (2019) Gene expression profile in patients with Gaucher disease indicates activation of inflammatory processes. Sci Rep 9:1–14. 10.1038/S41598-019-42584-1

Lukens JR, Gross JM, Calabrese C, et al (2014) Critical role for inflammasome-independent IL-1β production in osteomyelitis. Proc Natl Acad Sci U S A 111:1066–1071. 10.1073/PNAS.1318688111

Lull ME, Block ML (2010) Microglial activation and chronic neurodegeneration. Neurotherapeutics. 7:354–365. 10.1016/j.nurt.2010.05.014

Mata-Martínez E, Díaz-Muñoz M, Vázquez-Cuevas FG (2022) Glial cells and brain diseases: Inflammasomes as relevant pathological entities. Front Cell Neurosci 16:929529. 10.3389/FNCEL.2022.929529

Mink JW, Augustine EF, Adams HR, et al (2013) Classification and natural history of the neuronal ceroid lipofuscinoses. J Child Neurol 28:1101–1105. 10.1177/0883073813494268

Minkiewicz J, de Rivero Vaccari JP, Keane RW (2013) Human astrocytes express a novel NLRP2 inflammasome. Glia 61:1113–1121. 10.1002/glia.22499

Naeem A, Prakash R, Kumari N, et al (2024) MCC950 reduces autophagy and improves cognitive function by inhibiting NLRP3-dependent neuroinflammation in a rat model of Alzheimer’s disease. Brain Behav Immun 116:70–84. 10.1016/J.BBI.2023.11.031

Nagar A, Rahman T, Harton JA (2021) The ASC Speck and NLRP3 Inflammasome Function Are Spatially and Temporally Distinct. Front Immunol 12:752482. 10.3389/FNCEL.2022.929529

Nazir S, Gadi I, Al-Dabet MM, et al (2017) Cytoprotective activated protein C averts Nlrp3 inflammasome-induced ischemia-reperfusion injury via mTORC1 inhibition. Blood 130:2664–2677. 10.1182/BLOOD-2017-05-782102

Nourbakhsh F, Read MI, Barreto GE, Sahebkar A (2021) Astrocytes and inflammasome: A possible crosstalk in neurological diseases. Curr Med Chem 28:4972–4994. 10.2174/0929867328666210301105422

Okada M, Matsuzawa A, Yoshimura A, Ichijo H (2014) The lysosome rupture-activated TAK1-JNK pathway regulates NLRP3 inflammasome activation. Journal of Biological Chemistry 289:32926–32936. 10.1074/jbc.M114.579961

Paik S, Kim JK, Silwal P, et al (2021) An update on the regulatory mechanisms of NLRP3 inflammasome activation. Cell Mol Immunol 18:1141–1160. 10.1038/S41423-021-00670-3

Panicker N, Sarkar S, Harischandra DS, et al (2019) Fyn kinase regulates misfolded α-synuclein uptake and NLRP3 inflammasome activation in microglia. Journal of Experimental Medicine 928:175091. 10.1084/jem.20182191

Perera AP, Fernando R, Shinde T, et al (2018) MCC950, a specific small molecule inhibitor of NLRP3 inflammasome attenuates colonic inflammation in spontaneous colitis mice. Scientific Reports 2018 8:1 8:1–15. 10.1038/s41598-018-26775-w

Perry VH, Teeling J (2013) Microglia and macrophages of the central nervous system: the contribution of microglia priming and systemic inflammation to chronic neurodegeneration. Semin Immunopathol 35:601. 10.1007/S00281-013-0382-8

Pfaffl MW (2001) A new mathematical model for relative quantification in real-time RT–PCR. Nucleic Acids Res 29:e45–e45. 10.1093/NAR/29.9.E45

Polleux F, Ghosh A (2002) The slice overlay assay: a versatile tool to study the influence of extracellular signals on neuronal development. Sci STKE 2002: pl9. 10.1126/STKE.2002.136.PL9

Rangaraju S, Dammer EB, Raza SA, et al (2018) Identification and therapeutic modulation of a pro-inflammatory subset of disease-associated-microglia in Alzheimer’s disease. Mol Neurodegener 13:1–15. 10.1186/S13024-018-0254-8

Reinke S, Linge M, Diebner HH, et al (2020) Non-canonical Caspase-1 signaling drives RIP2-dependent and TNF-α-mediated inflammation in vivo. Cell Rep 30:2501–2511.e5. 10.1016/j.celrep.2020.01.090

Sahillioʇlu AC, Özören N (2015) Artificial loading of ASC specks with cytosolic antigens. PLoS One 10:e0134912. 10.1371/JOURNAL.PONE.0134912

Samant RR, Standaert DG, Harms AS (2024) The emerging role of disease-associated microglia in Parkinson’s disease. Front Cell Neurosci 18:1476461. 10.3389/FNCEL.2024.1476461

Sánchez KE, Bhaskar K, Rosenberg GA (2022) Apoptosis-associated speck-like protein containing a CARD-mediated release of matrix metalloproteinase 10 stimulates a change in microglia phenotype. Front Mol Neurosci 15: 976108. 10.3389/FNMOL.2022.976108

Saresella M, La Rosa F, Piancone F, et al (2016) The NLRP3 and NLRP1 inflammasomes are activated in Alzheimer’s disease. Molecular Neurodegeneration 2016 11:1 11:1–14. 10.1186/s13024-016-0088-1

Schulz A, Ajayi T, Specchio N, et al (2018) Study of intraventricular Cerliponase Alfa for CLN2 disease. New England Journal of Medicine 378:1898–1907. 10.1056/NEJMOA1712649

Scott XO, Stephens ME, Desir MC, et al (2020) The inflammasome adaptor protein ASC in Mild Cognitive Impairment and Alzheimer’s disease. Int J Mol Sci 21:1–22. 10.3390/IJMS21134674

Sleat DE, Wiseman JA, El-Banna M, et al (2004) A mouse model of classical late-infantile neuronal ceroid lipofuscinosis based on targeted disruption of the CLN2 gene results in a loss of tripeptidyl-peptidase I activity and progressive neurodegeneration. J Neurosci 24:9117–9126. 10.1523/JNEUROSCI.2729-04.2004

Van Opdenbosch N, Gurung P, Vande Walle L, et al (2014) Activation of the NLRP1b inflammasome independently of ASC-mediated caspase-1 autoproteolysis and speck formation. Nature Communications 2014 5:1 5:1–14. 10.1038/ncomms4209

Venegas C, Kumar S, Franklin BS, et al (2017) Microglia-derived ASC specks cross-seed amyloid-β in Alzheimer’s disease. Nature 552:355–361. 10.1038/NATURE25158

Ward R, Li W, Abdul Y, et al (2019) NLRP3 inflammasome inhibition with MCC950 improves diabetes-mediated cognitive impairment and vasoneuronal remodeling after ischemia. Pharmacol Res 142:237–250. 10.1016/J.PHRS.2019.01.035

White CS, Lawrence CB, Brough D, Rivers-Auty J (2017) Inflammasomes as therapeutic targets for Alzheimer’s disease. Brain Pathology 27:121–245. 10.1111/bpa.12478

Willingham SB, Bergstralh DT, O’Connor W, et al (2007) Microbial Pathogen-Induced Necrotic Cell Death Mediated by the Inflammasome Components CIAS1/Cryopyrin/NLRP3 and ASC. Cell Host Microbe 2:147–159. 10.1016/j.chom.2007.07.009

Xiong J, Kielian T (2013) Microglia in juvenile neuronal ceroid lipofuscinosis are primed toward a pro-inflammatory phenotype. J Neurochem 127:245–258. 10.1111/jnc.12385

Zengeler KE, Hollis A, Deutsch TCJ, et al (2025) Inflammasome signaling in astrocytes modulates hippocampal plasticity. Immunity 58:1519–1535.e11. 10.1016/j.immuni.2025.04.007

Zhang J, Pei L, Zang D, et al (2020) Gender Differences of NLRP1 Inflammasome in Mouse Model of Alzheimer’s Disease. Front Aging Neurosci 12:512097. 10.3389/FNAGI.2020.512097

