## Supplemental Figures for "The role of the NLRP3 inflammasome in a CLN2 mouse model of neuroinflammation"

### Supplemental Figure 1

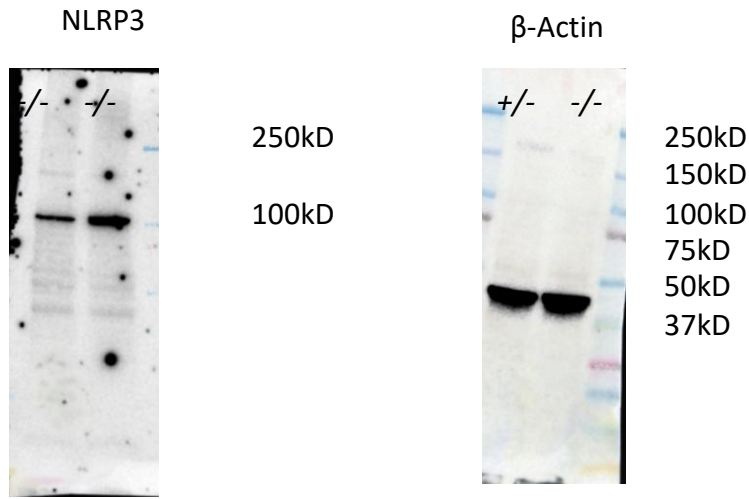

**Supplemental Figure 1:** Raw image of western blots shown and described in Figure 4A.

### Supplemental Figure 2

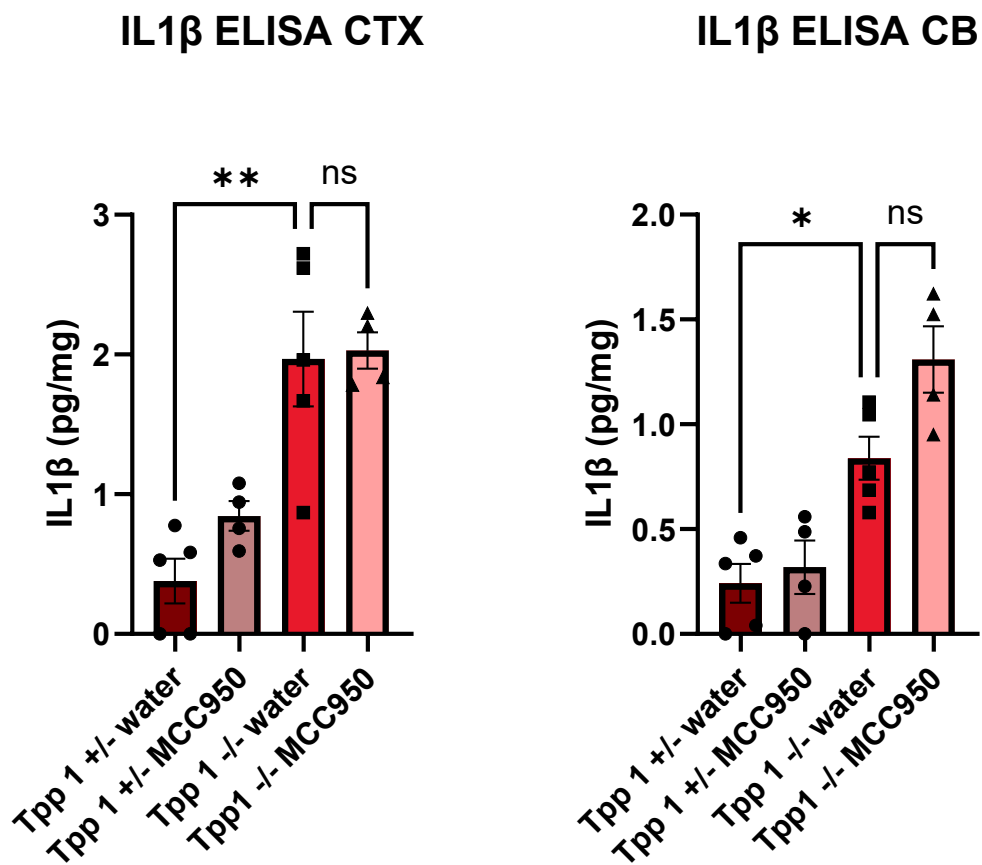

**Supplemental Figure 2:** MCC950 treatment does not impact IL1β protein levels. MCC950 was administered to *Tpp1* $-/-$ mice for the duration of 2-months in their drinking water and compared to control shams after sacrifice at 4-months of age, followed by an IL1β ELISA. IL1β concentration was not significantly different between *Tpp1* $-/-$ mice given control water and *Tpp1* $-/-$ mice provided MCC950 in either brain region.
